# Does science drive species naming, or can species naming drive science? A perspective from plant-feeding arthropods

**DOI:** 10.1101/2022.06.28.497962

**Authors:** Julia J. Mlynarek, Chloe Cull, Amy L. Parachnowitsch, Jess L. Vickruck, Stephen B. Heard

## Abstract

How do researchers choose their study species? Some choices are based on ecological or economic importance, some on ease of study, some on tradition – but could the name of a species influence researcher decisions? We asked whether phytophagous arthropod species named after their host plants were more likely to be assayed for host-associated genetic differentiation (or ‘HAD’; the evolution of cryptic, genetically isolated host specialists within an apparently more generalist lineage). We chose 30 arthropod species (from a Google Scholar search) for which a HAD hypothesis has been tested. We traced the etymologies of species names in the 30 corresponding genera, and asked whether HAD tests were more frequent among species whose etymologies were based on host-plant names (e.g., *Eurosta solidaginis*) vs. those with other etymologies (e.g., *Eurosta cribata*). Species with host-derived etymologies were more likely to feature in studies of HAD than those with other etymologies. We speculate that the etymology of a scientific name can draw a researcher’s attention to aspects of life-history and thus influence the direction of our scientific gaze.

## Introduction

Why do we study *this* species, and not *that* one? Some species are more charismatic; some are more fundable; some are more economically important; some are more abundant; some are found closer to home. The result is ecological and taxonomic bias in scientific attention. Some such biases are well known: for instance, arthropods are dramatically understudied compared to birds and mammals. Others are likely to be more subtle. These biases shape our understanding of the natural world, but seldom receive much explicit attention (although see Westoby 2002, Dietrich et al. 2020 for some discussion of how scientists should select study organisms, and how they actually do).

Could the names of species drive taxonomic bias in scientific attention? Since the mid-18^th^ century, scientists have used the Linnaean system to give each species a formal, Latinized name at the time of its scientific description. The bulk of species names follow a few etymological themes. Many species names refer to a species’ morphology, behaviour, geographical occurrence, or habitat, or are based on the name of a person (Figueiredo & Smith, 2010, Heard 2020, Mammalo et al. 2022, Poulin et al. 2022). However, there are few restrictions on how names can be formed or applied (even an arbitrary combination of letters can be a valid species name; Ride et al. 1999). Naming is thus an entirely creative act. Because humans are deeply interested in names, we wondered if the etymology of a species’ name might influence the kind of scientific attention later paid to it.

We investigated associations between species-name etymology and scientific study in the plant-feeding (or ‘phytophagous’) arthropods (insects and mites). The phytophagous arthropods are extraordinarily diverse, many are narrowly host-specific (Forister et al. 2015), and there is strong evidence that host specialization has been important in driving insect diversification (Matsubayishi et al. 2010, Nosil 2012, Forbes et al. 2017). As a result, evolutionary ecologists have frequently asked whether an apparently oligophagous or polyphagous plant-feeding arthropod (one that feeds on a few or many species of plants, respectively) might actually represent a complex of genetically distinct host forms each specializing on a single host species (e.g., Stireman et al. 2005, Scheffer and Hawthorne 2007, Mlynarek and Heard 2018). Such host forms are the result of “host-associated differentiation”, or HAD, an evolutionary process that disrupts random mating and allows independent adaptation of each resulting host form to its own host plant. The gallmaking tephritid fly *Eurosta solidaginis* is an excellent and well-studied example: what was once thought to be an oligophagous species attacking several goldenrod species is now understood to comprise genetically and ecologically distinct races specialized on *Solidago altissima* and *Solidago gigantea* (Abrahamson and Weis 1997), with further genetic structure associated with habitat in the west (Craig and Itami 2011) and likely with the alternative host *Solidago rugosa* in the east (Moffat et al. 2019).

While *Eurosta solidaginis* is unusual in the depth and breadth of attention paid to it, the basic question – one generalist, or a complex of specialists following HAD – has been asked of many other arthropods. But with hundreds of thousands (likely millions) of phytophagous species that might be studied, which ones tend to be assayed for the occurrence of HAD? We asked whether such assays are more likely for species whose scientific names are based on the names of their host plants – as is true for *E. solidaginis*. We speculated that such names might suggest to researchers, either consciously or subconsciously, that host specialization is an interesting part of the species’ biology and therefore ought to be studied. We compiled a dataset of species-name etymologies for 30 arthropod species that have been assayed for the presence of HAD, and for all their known congeners. We found that HAD-assayed species are, indeed, disproportionately likely to be species that were named for their food plants. This appears to be an example of taxonomic etymology directing, or at least influencing, the scientific gaze.

## Methods

### Data gathering

To find phytophagous arthropod species for which the hypothesis of host-associated differentiation was tested, we performed a Google Scholar search for the terms “HAD insect phytophagous herbivorous host associated differentiation”. In Scholar, this returns papers including *all* those terms in title or body, but the search is not case sensitive. The search returned about 25,900 results. We chose the first 30 papers that constituted tests of the HAD hypothesis for a phytophagous insect or mite using more than one host plant, disregarding whether the paper’s results supported or refuted the hypothesis of HAD. However, we removed one genus from our list: *Euura* (Roininen et al. 1993) has complex and uncertain genus-level taxonomy (Liston et al 2017), making some of our subsequent data-gathering steps impossible. We replaced *Euura* with *Nemorimyza* since *Nemorimyza posticata* Meigen, 1830 (a leaf-mining agromyzid fly) was recently tested for HAD (Mlynarek and Heard 2018) but was not flagged in our search. The resulting 30 HAD-tested species are listed in Table 1.

**Table 1.**
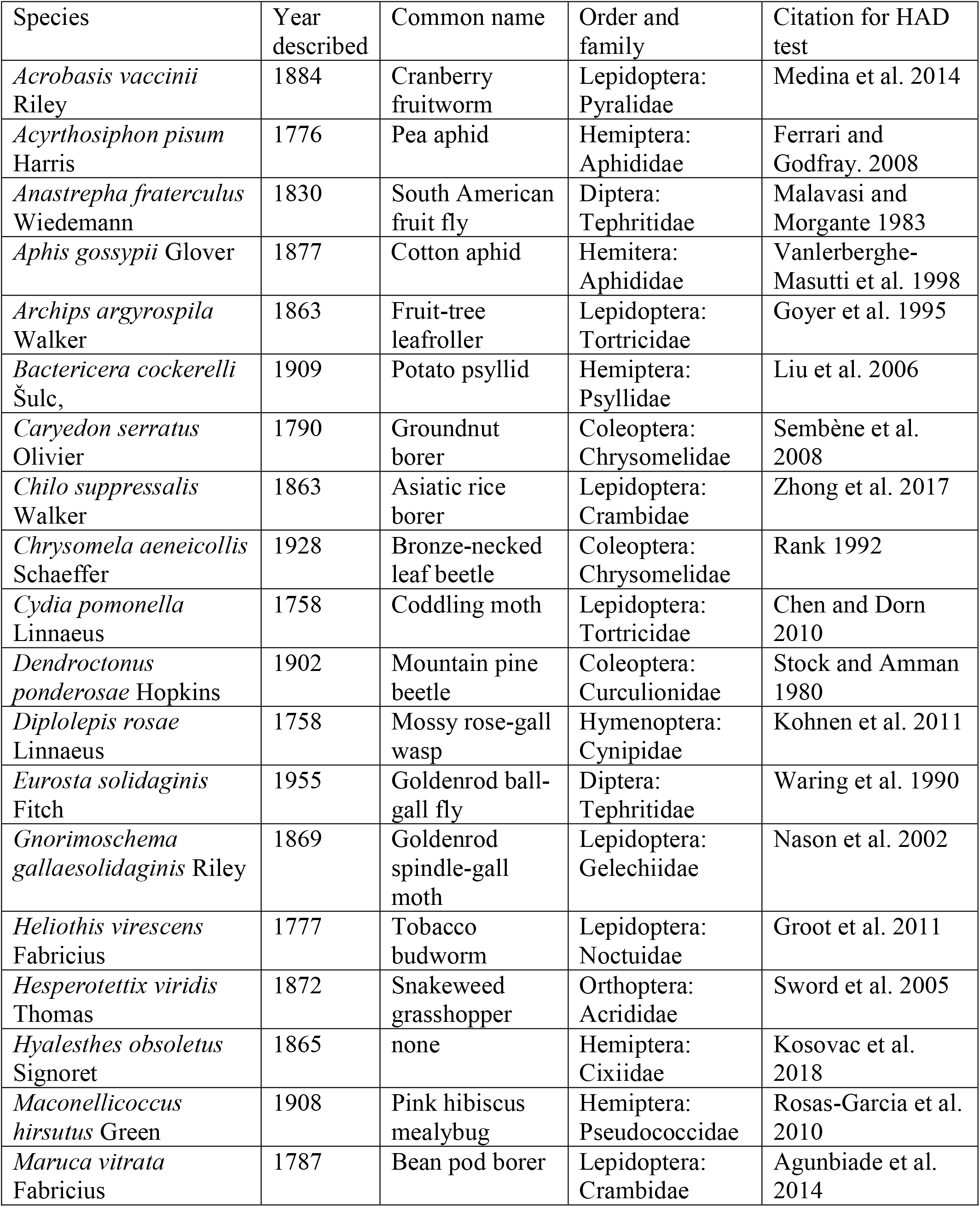
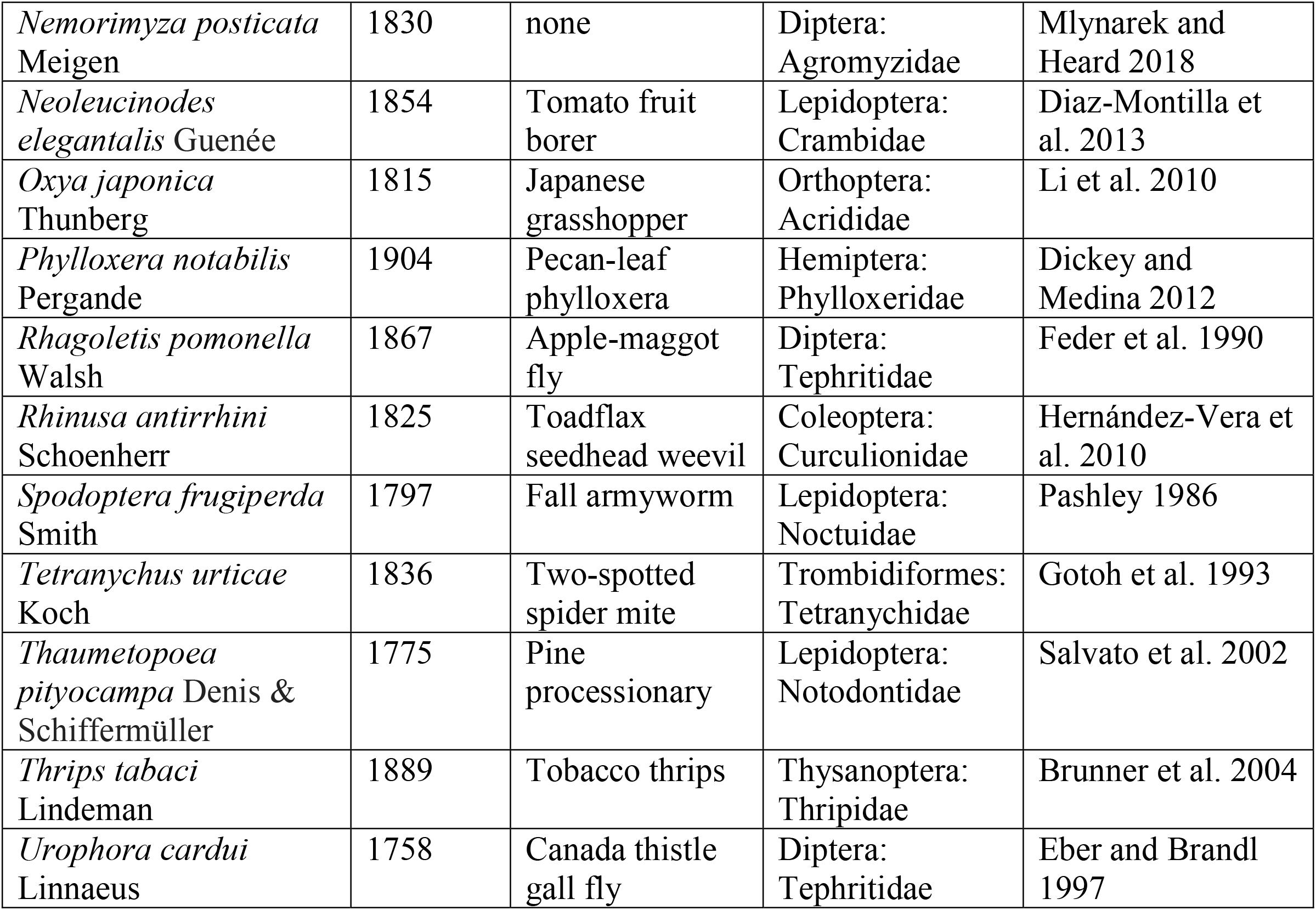
Phytophagous arthropod species, tested for the occurrence of host-associated genetic differentiation, identified by our search. The genus *Nemorimyza* replaced *Euura*, whose current phylogenetic resolution is too poor for our study.

| Species | Year described | Common name | Order and family | Citation for HAD test |
| --- | --- | --- | --- | --- |
| *Acrobasis vaccinii* Riley | 1884 | Cranberry fruitworm | Lepidoptera: Pyralidae | Medina et al. 2014 |
| *Acyrtosiphon pisum* Harris | 1776 | Pea aphid | Hemiptera: Aphididae | Ferrari and Godfray. 2008 |
| *Anastrepha fraterculus* Wiedemann | 1830 | South American fruit fly | Diptera: Tephritidae | Malavasi and Morgante 1983 |
| *Aphis gossypii* Glover | 1877 | Cotton aphid | Hemiptera: Aphididae | Vanlerberghe-Masutti et al. 1998 |
| *Archips argyrospila* Walker | 1863 | Fruit-tree leafroller | Lepidoptera: Tortricidae | Goyer et al. 1995 |
| *Bactericera cockerelli* Šulc, | 1909 | Potato psyllid | Hemiptera: Psyllidae | Liu et al. 2006 |
| *Caryedon serratus* Olivier | 1790 | Groundnut borer | Coleoptera: Chrysomelidae | Sembène et al. 2008 |
| *Chilo suppressalis* Walker | 1863 | Asiatic rice borer | Lepidoptera: Crambidae | Zhong et al. 2017 |
| *Chrysomela aeneicollis* Schaeffer | 1928 | Bronze-necked leaf beetle | Coleoptera: Chrysomelidae | Rank 1992 |
| *Cydia pomonella* Linnaeus | 1758 | Coddling moth | Lepidoptera: Tortricidae | Chen and Dorn 2010 |
| *Dendroctonus ponderosae* Hopkins | 1902 | Mountain pine beetle | Coleoptera: Curculionidae | Stock and Amman 1980 |
| *Diplolepis rosae* Linnaeus | 1758 | Mossy rose-gall wasp | Hymenoptera: Cynipidae | Kohnen et al. 2011 |
| *Eurosta solidaginis* Fitch | 1955 | Goldenrod ball-gall fly | Diptera: Tephritidae | Waring et al. 1990 |
| *Gnorimoschema gallaesolidaginis* Riley | 1869 | Goldenrod spindle-gall moth | Lepidoptera: Gelechiidae | Nason et al. 2002 |
| *Heliothis virescens* Fabricius | 1777 | Tobacco budworm | Lepidoptera: Noctuidae | Groot et al. 2011 |
| *Hesperotettix viridis* Thomas | 1872 | Snakeweed grasshopper | Orthoptera: Acrididae | Sword et al. 2005 |
| *Hyalesthes obsoletus* Signoret | 1865 | none | Hemiptera: Cixiidae | Kosovac et al. 2018 |
| *Maconellicoccus hirsutus* Green | 1908 | Pink hibiscus mealybug | Hemiptera: Pseudococcidae | Rosas-Garcia et al. 2010 |
| *Maruca vitrata* Fabricius | 1787 | Bean pod borer | Lepidoptera: Crambidae | Agunbiade et al. 2014 |
| *Nemorimyza posticata*<br>Meigen | 1830 | none | Diptera:<br>Agromyzidae | Mlynarek and<br>Heard 2018 |
| *Neoleucinodes*<br>*elegantalis* Guenée | 1854 | Tomato fruit<br>borer | Lepidoptera:<br>Crambidae | Diaz-Montilla et<br>al. 2013 |
| *Oxya japonica*<br>Thunberg | 1815 | Japanese<br>grasshopper | Orthoptera:<br>Acrididae | Li et al. 2010 |
| *Phylloxera notabilis*<br>Pergande | 1904 | Pecan-leaf<br>phylloxera | Hemiptera:<br>Phylloxeridae | Dickey and<br>Medina 2012 |
| *Rhagoletis pomonella*<br>Walsh | 1867 | Apple-maggot<br>fly | Diptera:<br>Tephritidae | Feder et al. 1990 |
| *Rhinusa antirrhini*<br>Schoenherr | 1825 | Toadflax<br>seedhead weevil | Coleoptera:<br>Curculionidae | Hernández-Vera et<br>al. 2010 |
| *Spodoptera frugiperda*<br>Smith | 1797 | Fall armyworm | Lepidoptera:<br>Noctuidae | Pashley 1986 |
| *Tetranychus urticae*<br>Koch | 1836 | Two-spotted<br>spider mite | Trombidiformes:<br>Tetranychidae | Gotoh et al. 1993 |
| *Thaumetopoea*<br>*pityocampa* Denis &<br>Schifferrmüller | 1775 | Pine<br>processionary | Lepidoptera:<br>Notodontidae | Salvato et al. 2002 |
| *Thrips tabaci*<br>Lindeman | 1889 | Tobacco thrips | Thysanoptera:<br>Thripidae | Brunner et al. 2004 |
| *Urophora cardui*<br>Linnaeus | 1758 | Canada thistle<br>gall fly | Diptera:<br>Tephritidae | Eber and Brandl<br>1997 |

For these species there are two possibilities. The species could have been described and named *after*, and perhaps even *as a result of*, a study documenting HAD; or the species might have been described and named *before* any such study was done. Only in the latter case could the etymology of the species’ name have influenced the researchers’ decision to study it, so we recorded both the year of taxonomic description and the year of HAD testing. This also allowed us to test for changes through time in naming practices.

We took our list of 30 focal HAD-tested species and compiled lists of all currently recognized species in each of their genera, using online resources and also by directly asking taxonomists with relevant expertise. There were 2,739 species distributed across the 30 genera. We then determined the etymology of the specific epithets. Where possible, we based our determination on the original species descriptions. When these descriptions did not include etymologies, or when they could not be located, we inferred the etymology from the linguistic formation (Latin or other root or suffix) of the name. For example, we inferred that a name ending in *-ensis* refers to a place of origin or distribution, whether or not this was explicitly indicated in the species description. We classified etymologies into seven categories: host, behavior, habitat, morphology, place, person (eponymy), or other. The “other” category was assigned either when we could not determine a specific epithet’s etymology, or when the etymology didn’t fit any of the other six categories. We included in the “other” category two few names based on host common names or cultural references: *Aphis sumire*, where “sumire” is a girls’ name meaning violet, and violet is the host; and *Phylloxera kunugi*, where “kunugi” is a Japanese common name for the oak species *Quercus acutissima*. We reasoned that although these etymologies are ultimately based on hosts, that connection would be inapparent to most of the global research community.

### Statistical analysis

We began by asking whether the distribution of etymologies differed among genera. This involves a G-test of independence applied to a 7 × 30 contingency table (7 etymological categories, 30 genera). We then used logistic regression to test the hypothesis that the proportion of species named for their host plant changed with year of description. In this analysis the response variable was a binary value indicating whether or not the arthropod species was named after its host.

To test our hypothesis that species-name etymology influenced likelihood of study, we asked whether those species that have been assayed for HAD were disproportionately named after their host plants, compared with all members of their genera. This is achieved using a G-test of independence to compare two proportions (in a 2 × 2 contingency table): first, the proportion of the 30 HAD-tested species with host-plant-based names, and second, the proportion of the 2,709 congeners with host-plant-based names. While a few of these congeners may themselves have been subjected to HAD testing, we did not attempt to separate such species out. This makes our G-test somewhat conservative. We repeated this G-test on two subsets of our data: first, we omitted *Aphis* (the largest genus, and something of an outlier in naming practice with by far the most host-plant-derived names); and second, we omitted *Tetranychus* (the only non-insect arthropod genus in our dataset).

All the analyses were performed in R using the packages *DescTools* (Signorell et al. 2021) and *vegan* 2.5-7 (Oksanen et al. 2020).

## Results

In our dataset, for all species with published HAD studies, the naming preceded the HAD test (Table 1, compare 2^nd^ and last columns).

There was substantial variation among genera in the breakdown of species-name etymologies (*G* = 1052, df = 174, *P* < 2.2 × 10^−16^; Table 2). Morphology (895 species), host (646 species), person (423 species) and place (300 species) were the most popular origins of names given to species. Names referring to host plants are most common (57%) in *Aphis*, which is the only genus to exceed the proportion of host-plant names for our HAD species; Figure 1). Three genera (*Hesperotettix, Hyalesthes*, and *Maruca*) have no species named after host plants, and for *Eurosta*, only the focal *E. solidaginis* has such a name. For the remaining genera, between 4 and 37% of species were named after plants. Species were significantly more likely to be named after their hosts if described earlier (*z* = 3.93, *P* < 0.001. However, the effect size was only moderate: between 1760 and 2020, the modelled fraction of insects named after host plants declined from about 36% to 21% (Figure 2).

**Table 2.** Etymologies of species epithets in 30 genera of plant-feeding arthropods. “Other” includes etymologies that are unclear as well as those not fitting the remaining categories.

| Genus | behaviour | habitat | morphology | person | place | other | host | All non-host categories | TOTAL |
| --- | --- | --- | --- | --- | --- | --- | --- | --- | --- |
| Acrobasis | 0 | 9 | 56 | 18 | 15 | 18 | 23 | 116 | 139 |
| Acyrtosiphon | 0 | 6 | 15 | 17 | 9 | 9 | 35 | 56 | 91 |
| Anastrepha | 0 | 8 | 140 | 75 | 23 | 43 | 10 | 289 | 299 |
| Aphis | 12 | 12 | 57 | 84 | 42 | 54 | 340 | 261 | 601 |
| Archips | 1 | 6 | 50 | 17 | 12 | 20 | 12 | 106 | 118 |
| Bactericera | 0 | 5 | 47 | 19 | 15 | 9 | 28 | 95 | 123 |
| Caryedon | 0 | 0 | 2 | 1 | 3 | 2 | 3 | 8 | 11 |
| Chilo | 0 | 2 | 23 | 3 | 10 | 15 | 5 | 53 | 58 |
| Chrysomela | 0 | 0 | 19 | 4 | 5 | 4 | 6 | 32 | 38 |
| Cydia | 1 | 10 | 68 | 22 | 13 | 79 | 24 | 193 | 217 |
| Dendroctonus | 0 | 0 | 11 | 1 | 2 | 1 | 3 | 15 | 18 |
| Diplolepis | 0 | 3 | 29 | 7 | 5 | 5 | 2 | 49 | 51 |
| Eurosta | 0 | 0 | 5 | 0 | 1 | 0 | 1 | 6 | 7 |
| Gnorimoschema | 0 | 6 | 47 | 23 | 7 | 22 | 10 | 105 | 115 |
| Heliothis | 1 | 2 | 27 | 4 | 4 | 7 | 1 | 45 | 46 |
| Hesperotettix | 0 | 0 | 3 | 1 | 4 | 1 | 0 | 9 | 9 |
| Hyalesthes | 0 | 0 | 12 | 3 | 12 | 4 | 0 | 31 | 31 |
| Maconellicoccus | 0 | 0 | 3 | 0 | 4 | 0 | 1 | 7 | 8 |
| Maruca | 0 | 0 | 3 | 0 | 1 | 0 | 0 | 4 | 4 |
| Nemorimyza | 0 | 0 | 3 | 0 | 2 | 0 | 1 | 5 | 6 |
| Neoleucinodes | 0 | 2 | 5 | 0 | 0 | 0 | 1 | 7 | 8 |
| Oxya | 1 | 2 | 25 | 2 | 16 | 3 | 0 | 49 | 49 |
| Phylloxera | 2 | 0 | 26 | 4 | 4 | 2 | 17 | 38 | 55 |
| Rhagoletis | 0 | 1 | 23 | 14 | 7 | 6 | 13 | 51 | 64 |
| Rhinusa | 0 | 0 | 11 | 4 | 2 | 1 | 5 | 18 | 23 |
| Spodoptera | 0 | 1 | 19 | 1 | 4 | 12 | 0 | 37 | 37 |
| Tetranychus | 0 | 1 | 21 | 38 | 33 | 14 | 41 | 107 | 148 |
| Thaumetopoea | 2 | 0 | 0 | 2 | 2 | 1 | 2 | 7 | 9 |
| Thrips | 0 | 7 | 118 | 42 | 31 | 29 | 57 | 227 | 284 |
| Urophora | 0 | 1 | 27 | 16 | 12 | 10 | 6 | 66 | 72 |

**Figure 1.**
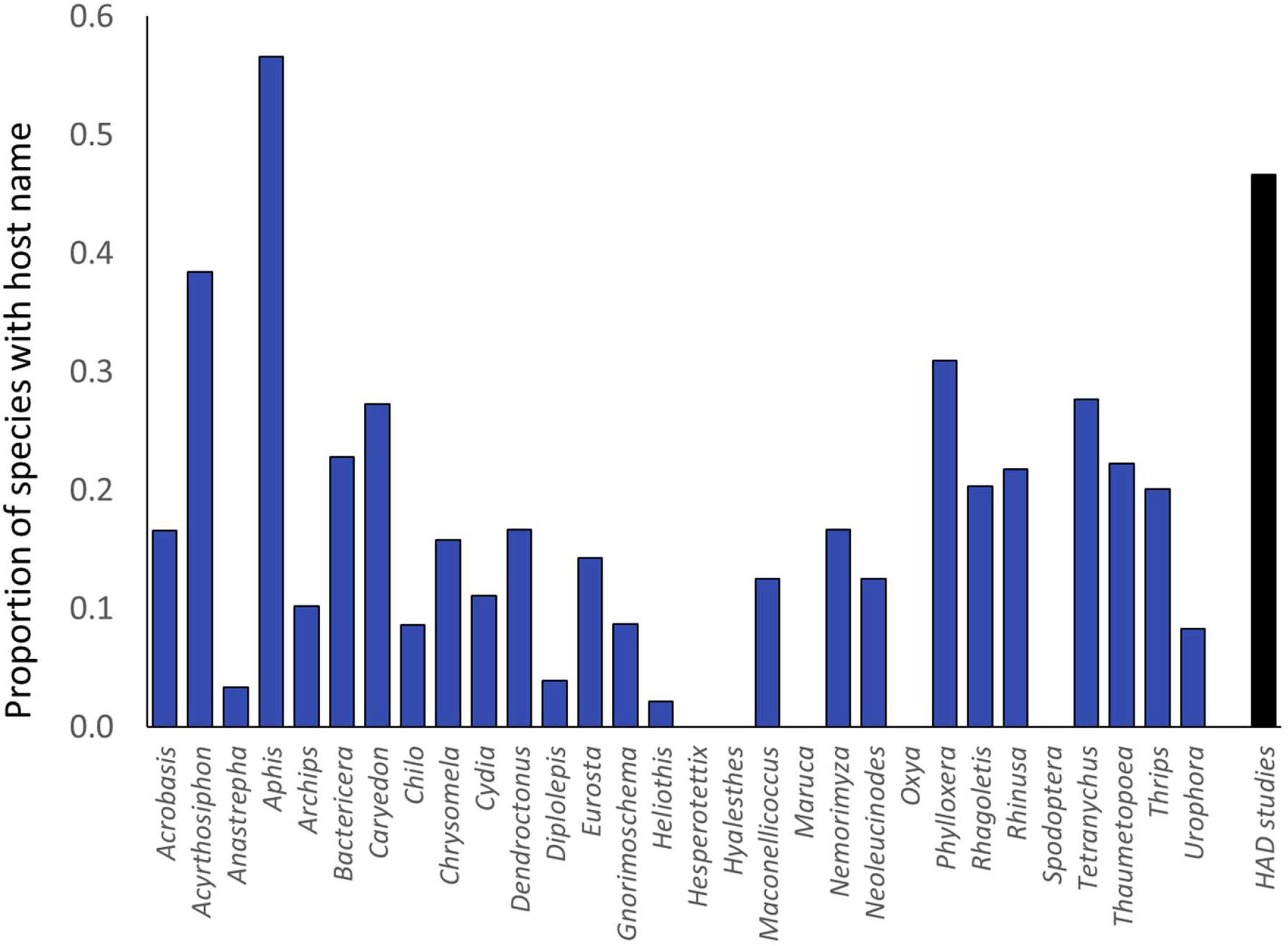
Proportion of species named for host plants across the 30 focal genera, compared with the proportion for the 30 focal HAD-tested species (black bar at right). The plotted proportions for each genus exclude the focal species.

**Figure 2.**
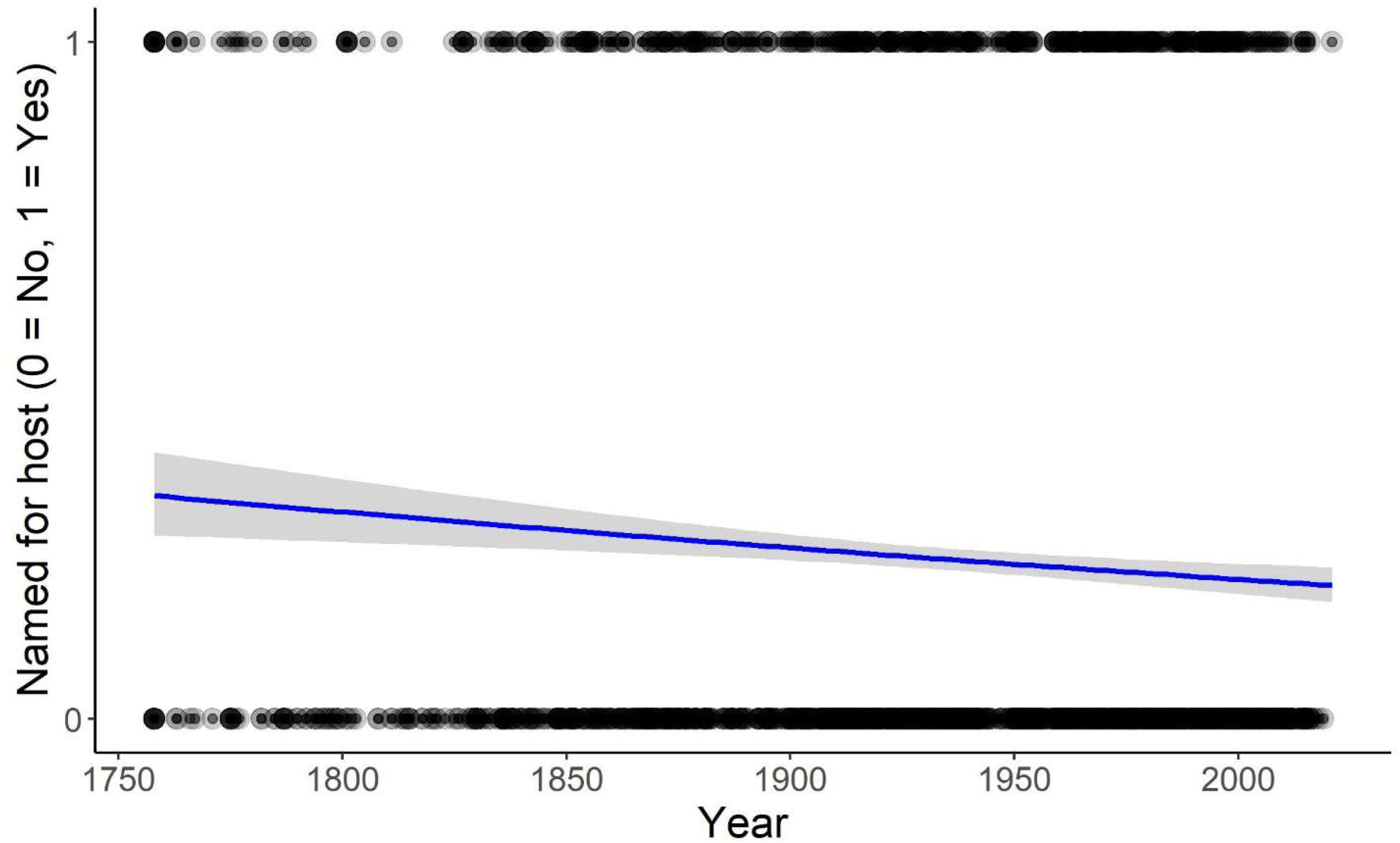
Arthropod species that were named for their plant host, or otherwise, organized by their year of description. Blue line represents the probability curve generated from the logistic regression.

Of our 30 focal HAD-tested species, 14 (47%) were named after a host plant. A significantly smaller proportion of the 2,709 congeners (632 species, or 23%) had this etymology (*G* = 7.67, df = 1, *P* = 0.0056). This pattern remains robust if we omit from the analysis either *Aphis* (*G* = 15.8, df = 1, *P* = 0.00007) or *Tetranychus* (*G* = 6.49, df = 1, *P* = 0.01).

## Discussion

There are far more species on Earth than can possibly be studied; and even among species that do attract the scientific gaze, some are studied far more than others. Using data for phytophagous arthropods, we asked whether the etymology of a species name –in particular, whether it is named after a host plant – might influence the kind of scientific attention that species receives.

We found striking and significant variation among genera in the use of different etymological categories in constructing species names. This included strong variation in the proportion of names based on plant hosts (from just one species in *Eurosta* to over half of the speciose genus *Aphis*). Poulin et al. (2022) documented a similar pattern (but at coarser phylogenetic scale) among parasitic worms, with strong variation among classes and phyla in naming etymologies. This kind of variation is not unexpected (Poulin et al. 2022), given that different taxonomists are involved with naming in different clades. We also found a tendency, significant but weak, for names based on host plants to have become less common over time. Mammola et al. (2022) found a similar but stronger effect for ecologically informative names among spiders, although Poulin et al. (2022) found no such trend for parasitic worms. These patterns underscore the fundamentally creative nature of the naming act. The assignment of a specific epithet to a newly described species is ultimately an arbitrary and creative decision (Heard 2020) that might reflect the biology of the named species, but can also (or instead) reflect the culture and personalities of the taxonomists assigning the name. And yet naming is potentially important, because humans are extraordinarily interested in names and may perceive information in names even if that perception isn’t intended by the namer.

The etymological basis of a species name was significantly associated with the likelihood of that species being studied as a potential example of host-associated differentiation (HAD). In particular, species with names based on those of their host plants were strongly overrepresented in HAD studies. While our analysis cannot break this pattern down to the level of individual genera, we were able to rule out the possibility of its being driven by two potential outliers: *Aphis*, a diverse genus for which naming practices have been somewhat unusual (with far more host-species-derived names than any other genus), and *Tetranychus*, the only non-insect arthropod in our dataset. Our results suggest that decisions made by the taxonomists describing species, often decades or centuries earlier, are shaping the attention evolutionary ecologists pay to taxa now. Because the frequency of host-plant names varies across insect taxa, this effect could be driving a taxonomic bias in HAD studies – and as a result, in the detection of HAD – across arthropod lineages.

Why might researchers disproportionately study host-plant-named arthropods? We have two speculations. First, researchers may notice the name and be subconsciously inspired to ask questions about host specialization. One of us (SBH) first came to the study of HAD by noticing moth galls on stems of *Solidago altissma* and *Solidago gigantea*, and wondering if they represented a pair of host specialists. It’s certainly possible that learning the gallmaker’s name – *Gnorimoschema gallaesolidaginis* – played some role in feeding this wonder. Second, perhaps pest species are more likely to be studied to answer evolutionary ecology questions, in part because their economic impact makes the work more fundable. If these pests are more likely named after the host plant they impact, then our HAD-naming pattern could arise. In this vein, it is interesting that among our 30 focal genera, the highest proportion of host-plant names occurred in *Aphis* – a genus including many agricultural pests.

One might wonder whether species names might bias not just the *occurrence* of HAD studies, but also their outcome. This could be true if insects named for their host plants are more likely to be strict specialists. Under this scenario, the same species researchers are most likely to assay for HAD are the ones most likely to reveal that HAD has occurred. The result would be exaggerated estimates of the frequency of HAD among plant-feeding insects, and so this is a disquieting possibility. We are not, unfortunately, in a position to shed much more light on this. It seems plausible that oligophagous species are more likely to be named for their hosts than broad generalists, but it does not necessarily follow that these oligophages are more likely to in fact represent complexes of cryptic host specialists. Ideally, we would compare the frequency with which assays for HAD actually reveal its occurrence between taxa named for plant hosts and those named in other ways. However, given the file drawer effect (studies that don’t find HAD may be less likely to be published) and given the relatively small number of insects for which powerful tests of HAD are available, this will have to remain a goal for the (distant) future.

Might there be other ways in which naming influences later scientific attention to species? We are unaware of data bearing on this, but we suspect so. At the most obvious level, taxa without formal names are unlikely to be studied. This might well be true even for taxa that have been identified as distinct and thus nameable lineages, but have not yet been formally described and named (e.g., the divergent habitat- and host-associated clades within *Nemorimyza posticata* identified by Mlynarek and Heard 2018). Thinking further along these lines, we wonder whether species epithets that are very long, difficult to spell, or difficult to pronounce might reduce the scientific attention paid to their bearers. This hypothesis should be testable with current data, or we can wait to assess the future literature corpus devoted to the recently described myxobacterium *Myxococcus llanfairpwllgwyngyllgogerychwyrndrobwllllantysiliogogogochensis* (Chambers et al. 2020).

Our study provides yet more evidence (if it was needed) that the common stereotype of science as objective and fully rational is ill-founded. Scientists study a non-random subset of the world, and ask a non-random subset of questions. We have provided evidence that these two dimensions of non-randomness may be interrelated. Scientists, like novelists and songwriters, are often asked where they get their ideas. It would appear that sometimes the answer lies in names.

## Acknowledgements

We are grateful to members of the Population Ecology and Evolution Research Group, University of New Brunswick, for comments on this work and to Robert Poulin for an advance look at some related research. This work was funded in part by grant support from the Natural Sciences and Engineering Research Council (Canada; Discovery Grant RGPIN-2015-04418 to SBH).

## References

Abrahamson WG, Weis AE. 1997 Evolutionary Ecology across Three Trophic Levels: Goldenrods, Gallmakers, and Natural Enemies. Princeton, NJ: Princeton University Press.

Agunbiade TA, Coates BS, Datinon B, Djouaka R, Sun W, Tamò M, Pittendrigh BR. 2014 Genetic differentiation among Maruca vitrata F. (Lepidoptera: Crambidae) populations on cultivated cowpea and wild host plants: Implications for insect resistance management and biological control strategies. PLoS ONE 9. (doi:10.1371/journal.pone.0092072)

Brunner PC, Chatzivassiliou EK, Katis NI, Frey JE. 2004 Host-associated genetic differentiation in Thrips tabaci (Insecta; Thysanoptera), as determined from mtDNA sequence data. Heredity 93, 364–370. (doi:10.1038/sj.hdy.6800512)

Chambers J, Sparks N, Sydney N, Livingstone PG, Cookson AR, Whitworth DE. 2020 Comparative Genomics and Pan-Genomics of the Myxococcaceae, including a Description of Five Novel Species: Myxococcus eversor sp. nov., Myxococcus llanfairpwllgwyngyllgogerychwyrndrobwllllantysiliogogogochensis sp. nov., Myxococcus vastator sp. nov., Pyxidicoccus caerfyrddinensis sp. nov., and Pyxidicoccus trucidator sp. nov. Genome Biol Evol 12, 2289–2302. (doi:10.1093/gbe/evaa212)

Chen MH, Dorn S. 2010 Microsatellites reveal genetic differentiation among populations in an insect species with high genetic variability in dispersal, the codling moth, Cydia pomonella (L.) (Lepidoptera: Tortricidae). Bulletin of Entomological Research 100, 75–85. (doi:10.1017/S0007485309006786)

Craig TP, Itami JK. 2011 Divergence of Eurosta solidaginis in response to host plant variation and natural enemies. Evolution 65, 802–817. (doi:10.1111/j.1558-5646.2010.01167.x)

Díaz-Montilla AE, Suárez-Baron HG, Gallego-Sánchez G, Saldamando-Benjumea CI, Tohme J. 2013 Geographic differentiation of Colombian Neoleucinodes elegantalis (Lepidoptera: Crambidae) haplotypes: Evidence for Solanaceae host plant association and Holdridge life zones for genetic differentiation. Annals of the Entomological Society of America 106, 586–597. (doi:10.1603/AN12111)

Dickey AM, Medina RF. 2012 Host-associated genetic differentiation in pecan leaf phylloxera. Entomologia Experimentalis et Applicata 143, 127–137. (doi:10.1111/j.1570-7458.2012.01250.x)

Dietrich MR, Ankeny RA, Crowe N, Green S, Leonelli S. 2020 How to choose your research organism. Studies in History and Philosophy of Science Part C :Studies in History and Philosophy of Biological and Biomedical Sciences 80. (doi:10.1016/j.shpsc.2019.101227)

Eber S, Brandl R. 1997 Genetic differentiation of the tephritid fly Urophora cardui in Europe as evidence for its biogeographical history. Molecular Ecology 6, 651–660. (doi:10.1046/j.1365-294X.1997.00236.x)

Feder JL, Chilcote CA, Bush GL. 1990 The geographic pattern of genetic differentiation between host associated populations of Rhagoletis pomonella (Diptera: Tephritidae) in the eastern United states and Canada. Evolution 44, 570–594. (doi:10.1111/j.1558-5646.1990.tb05939.x)

Ferrari J, Via S, Godfray HCJ. 2008 Population differentiation and genetic variation in performance on eight hosts in the pea aphid complex. Evolution 62, 2508–2524. (doi:10.1111/j.1558-5646.2008.00468.x)

Figueiredo E, Smith GF. 2010 What’s in a name: Epithets in Aloe L. (Asphodelaceae) and what to call the next new species. Bradleya 28, 79–102. (doi:10.25223/brad.n28.2010.a9)

Forbes AA, Devine SN, Hippee AC, Tvedte ES, Ward AKG, Widmayer HA, Wilson CJ. 2017 Revisiting the particular role of host shifts in initiating insect speciation. Evolution 71, 1126–1137. (doi:10.1111/evo.13164)

Forister ML et al. 2015 The global distribution of diet breadth in insect herbivores. Proceedings of the National Academy of Sciences of the United States of America 112, 442–447. (doi:10.1073/pnas.1423042112)

Gotoh T, Bruin J, Sabelis MW, Menken SBJ. 1993 Host race formation in Tetranychus urticae: genetic differentiation, host plant preference, and mate choice in a tomato and a cucumber strain. Entomologia Experimentalis et Applicata 68, 171–178. (doi:10.1111/j.1570-7458.1993.tb01700.x)

Goyer RA, Paine TD, Pashley DP, Lenhard GJ, Meeker JR, Hanlon CC. 1995 Geographic and host-associated differentiation in the fruittree leafroller (Lepidoptera: Tortricidae). Annals of the Entomological Society of America 88, 391–396. (doi:10.1093/aesa/88.4.391)

Groot AT, Classen A, Inglis O, Blanco CA, LO’Pez Jr. J, TÉran Vargas A, Schal C, Heckel DG, SchÖfl G. 2011 Genetic differentiation across North America in the generalist moth Heliothis virescens and the specialist H. subflexa. Molecular Ecology 20, 2676–2692. (doi:10.1111/j.1365-294X.2011.05129.x)

Heard SB. 2022 Charles Darwin’s Barnacle and David Bowie’s Spider. New Haven: Yale University Press. See https://yalebooks.yale.edu/9780300238280/charles-darwins-barnacle-and-david-bowies-spider.

Hernández-Vera G, Mitrović M, Jović J, Toševski I, Caldara R, Gassmann A, Emerson BC. 2010 Host-associated genetic differentiation in a seed parasitic weevil Rhinusa antirrhini (Coleptera: Curculionidae) revealed by mitochondrial and nuclear sequence data. Molecular Ecology 19, 2286–2300. (doi:10.1111/j.1365-294X.2010.04639.x)

Kohnen A, Wissemann V, Brandl R. 2011 No host-associated differentiation in the gall wasp Diplolepis rosae (Hymenoptera: Cynipidae) on three dog rose species. Biological Journal of the Linnean Society 102, 369–377. (doi:10.1111/j.1095-8312.2010.01582.x)

Kosovac A, Johannesen J, Krstić O, Mitrović M, Cvrković T, Toševski I, Jović J. 2018 Widespread plant specialization in the polyphagous planthopper Hyalesthes obsoletus (Cixiidae), a major vector of stolbur phytoplasma: Evidence of cryptic speciation. PLoS ONE 13. (doi:10.1371/journal.pone.0196969)

Li T, Geng Y-P, Zhong Y, Zhang M, Ren Z-M, Ma J, Guo Y-P, Ma E-B. 2010 Host-associated genetic differentiation in rice grasshopper, Oxya japonica, on wild vs. cultivated rice. Biochemical Systematics and Ecology 38, 958–963. (doi:10.1016/j.bse.2010.05.003)

Liston AD, Heibo E, Prous M, Vårdal H, Nyman T, Vikberg V. 2017 North European gall-inducing Euura sawflies (Hymenoptera, Tenthredinidae, Nematinae). Zootaxa 4302, 1–115. (doi:10.11646/zootaxa.4302.1.1)

Liu D, Trumble JT, Stouthamer R. 2006 Genetic differentiation between eastern populations and recent introductions of potato psyllid (Bactericera cockerelli) into western North America. Entomologia Experimentalis et Applicata 118, 177–183. (doi:10.1111/j.1570-7458.2006.00383.x)

Malavasi A, Morgante JS. 1983 Population genetics of Anastrepha fraterculus (Diptera, Tephritidae) in different hosts: Genetic differentiation and heterozygosity. Genetica 60, 207–211. (doi:10.1007/BF00122375)

Mammola S, Viel N, Amiar D, Mani A, Hervé C, Heard SB, Fontaneto D, Pétillon J. In press. Taxonomic practice, creativity, and fashion: What’s in a spider name? (doi:https://doi.org/10.1101/2022.02.06.479275)

Matsubayashi KW, Ohshima I, Nosil P. 2010 Ecological speciation in phytophagous insects. Entomologia Experimentalis et Applicata 134, 1–27. (doi:10.1111/j.1570-7458.2009.00916.x)

Medina RF, Szendrei Z, Harrison K, Isaacs R, Averill A, Malo EA, Rodriguez-Saona C. 2014 Exploring host-associated differentiation in the North American native cranberry fruitworm, Acrobasis vaccinii, from blueberries and cranberries. Entomologia Experimentalis et Applicata 150, 136–148. (doi:10.1111/eea.12143)

Mlynarek JJ, Heard SB. 2018 Strong and complex host- and habitat-associated genetic differentiation in an apparently polyphagous leaf mining insect. Biological Journal of the Linnean Society 125, 885–899. (doi:10.1093/biolinnean/bly166)

Moffat C, Takahashi M, Pease S, Brown J, Heard S, Abrahamson W. 2019 Are Eurosta solidaginis on Solidago rugosa a divergent host-associated race? Evolutionary Ecology 33. (doi:10.1007/s10682-018-9966-z)

Nason JD, Heard SB, Williams FR. 2002 Host-associated genetic differentiation in the goldenrod elliptical-gall moth, Gnorimoschema gallaesolidaginis (Lepidoptera: Gelechiidae). Evolution 56, 1475–1488. (doi:10.1111/j.0014-3820.2002.tb01459.x)

Nosil P. 2012 Ecological Speciation. Oxford, United Kingdom: Oxford University Press.

Oksanen J et al. 2022 vegan: Community Ecology Package. See https://CRAN.R-project.org/package=vegan.

Pashley DP. 1986 Host-associated genetic differentiation in fall armyworm (Lepidoptera: Noctuidae): A sibling species complex? Annals of the Entomological Society of America 79, 898–904. (doi:10.1093/aesa/79.6.898)

Poulin R, McDougall C, Presswell B. 2022 What’s in a name? Taxonomic and gender biases in the etymology of new species names. Proceedings of the Royal Society B: Biological Sciences 289, 20212708. (doi:10.1098/rspb.2021.2708)

Rank NE. 1992 A hierarchical analysis of genetic differentiation in a montane leaf beetle Chrysomela aeneicollis (Coleoptera: Chrysomelidae). Evolution 46, 1097–1111. (doi:10.1111/j.1558-5646.1992.tb00622.x)

Ride WDL, International Commission on Zoological Nomenclature, International Union of Biological Sciences. 1999 International Code of Zoological Nomenclature: Fourth Edition. London, United Kingdom: International Trust for Zoological Nomenclature, c/o Natural History Museum.

Roininen H, Vuorinen J, Tahvanainen J, Julkunen-Tiitto R. 1993 Host preference and allozyme differentiation in shoot galling sawfly, Euura atra. Evolution 47, 300–308. (doi:10.2307/2410137)

Rosas-García NM, Sarmiento-Benavides SL, Villegas-Mendoza JM, Hernández-Delgado S, Mayek-Pérez N. 2010 Genetic differentiation among Maconellicoccus hirsutus (Hemiptera: Pseudococcidae) populations living on different host plants. Environmental Entomology 39, 1043–1050. (doi:10.1603/EN09368)

Salvato P, Battisti A, Concato S, Masutti L, Patarnello T, Zane L. 2002 Genetic differentiation in the winter pine processionary moth (Thaumetopoea pityocampa - wilkinsoni complex), inferred by AFLP and mitochondrial DNA markers. Molecular Ecology 11, 2435–2444. (doi:10.1046/j.1365-294X.2002.01631.x)

Scheffer SJ, Hawthorne DJ. 2007 Molecular evidence of host-associated genetic divergence in the holly leafminer Phytomyza glabricola (Diptera: Agromyzidae): Apparent discordance among marker systems. Molecular Ecology 16, 2627–2637. (doi:10.1111/j.1365-294X.2007.03303.x)

Sembène M, Rasplus J-Y, Silvain J-F, Delobel A. 2008 Genetic differentiation in sympatric populations of the groundnut seed beetle Caryedon serratus (Coleoptera: Chrysomelidae): new insights from molecular and ecological data. International Journal of Tropical Insect Science 28, 168–177. (doi:10.1017/S1742758408094484)

Signorell A et al. 2022 DescTools: Tools for Descriptive Statistics. See https://CRAN.R-project.org/package=DescTools.

Stireman JO, Nason JD, Heard SB. 2005 Host-associated genetic differentiation in phytophagous insects: general phenomenon or isolated exceptions? Evidence from a goldenrod-insect community. Evolution 59, 2573–2587.

Stock MW, Amman GD. 1980 Genetic differentiation among mountain pine beetle populations from lodgepole pine and ponderosa pine in northeast Utah1. Annals of the Entomological Society of America 73, 472–478. (doi:10.1093/aesa/73.4.472)

Sword GA, Joern A, Senior LB. 2005 Host plant-associated genetic differentiation in the snakeweed grasshopper, Hesperotettix viridis (Orthoptera: Acrididae). Molecular Ecology 14, 2197–2205. (doi:10.1111/j.1365-294X.2005.02546.x)

Vanlerberghe-Masutti F, Chavigny P. 1998 Host-based genetic differentiation in the aphid Aphis gossypii Glover, evidenced from RAPD fingerprints. Molecular Ecology 7, 905–914. (doi:10.1046/j.1365-294x.1998.00421.x)

Waring GL, Abrahamson WG, Howard DJ. 1990 Genetic differentiation among host-associated populations of the gallmaker Eurosta solidaginis (Diptera: Tephritidae). Evolution 44, 1648–1655. (doi:10.1111/j.1558-5646.1990.tb03853.x)

Westoby M. 2002 Choosing species to study. Trends in Ecology & Evolution 17, 587. (doi:10.1016/S0169-5347(02)02634-4)

Zhong H, Li F, Chen J, Zhang J, Li F. 2017 Comparative transcriptome analysis reveals host-associated differentiation in Chilo suppressalis (Lepidoptera: Crambidae). Scientific Reports 7. (doi:10.1038/s41598-017-14137-x)

